# Demethylmenaquinone methyl transferase is a membrane domain-associated protein essential for menaquinone homeostasis in *Mycobacterium smegmatis*

**DOI:** 10.1101/452839

**Authors:** Julia Puffal, Jacob A. Mayfield, D. Branch Moody, Yasu S. Morita

## Abstract

The intracellular membrane domain (IMD) in mycobacteria is a spatially distinct region of the plasma membrane with diverse functions. Previous comparative proteomic analysis of the IMD suggested that menaquinone biosynthetic enzymes are associated with this domain. In the present study, we determined the subcellular site of these enzymes using sucrose density gradient fractionation. We found that the last two enzymes, the methyltransferase MenG, and the reductase MenJ, are associated with the IMD. MenA, the prenyltransferase that mediates the first membrane-associated step of the menaquinone biosynthesis, is associated with the conventional plasma membrane. For MenG, we additionally showed the polar enrichment of the fluorescent protein fusion colocalizing with an IMD marker protein *in situ*. To start dissecting the roles of IMD-associated enzymes, we further tested the physiological significance of MenG. The deletion of *menG* at the endogenous genomic loci was possible only when an extra copy of the gene was present, indicating that it is an essential gene in *M. smegmatis*. Using a tetracycline-inducible switch, we achieved gradual and partial depletion of MenG over three consecutive 24 hour subcultures. This partial MenG depletion resulted in progressive slowing of growth, which corroborated the observation that *menG* is an essential gene. Upon MenG depletion, there was a significant accumulation of MenG substrate, demethylmenaquinone, even though the cellular level of menaquinone, the reaction product, was unaffected. Furthermore, the growth retardation was coincided with a lower oxygen consumption rate and ATP accumulation. These results imply a previously unappreciated role of MenG in regulating menaquinone homeostasis within the complex spatial organization of mycobacterial plasma membrane.

## Introduction

*Mycobacterium smegmatis* has a complex membrane organization. In addition to the topologically distinct outer mycolyl and inner plasma membranes, the plasma membrane has a spatially distinct membrane domain known as the Intracellular Membrane Domain (IMD) (Hayashi et al., 2016; 2018). Experimentally, the IMD can be separated and purified from the conventional plasma membrane by sucrose density gradient fractionation of mycobacterial crude cell lysate (Morita et al., 2005). In this gradient fractionation, the IMD appears as vesicles of phospholipids without significant enrichment of cell wall components. In contrast, the conventional plasma membrane fraction contains both membrane phospholipids and cell wall components, suggesting that the conventional plasma membrane is tightly associated with the cell wall (designated as PM-CW). A more recent study revealed that the IMD is particularly enriched in the polar regions of the live actively growing cell, and associated with more than 300 proteins, among which are enzymes involved in cell envelope biosynthesis (Hayashi et al., 2016). Mycobacteria extend their cell envelope primarily from the polar region of the rod-shaped cell, and unlike other model bacteria such as *Escherichia coli* or *Bacillus subtilis*, the cylindrical part of the cell does not actively elongate (Aldridge et al., 2012; Thanky et al., 2007). Therefore, the polar IMD enrichment implies the strategic placement of membrane-bound enzymes that are involved in producing cell envelope biosynthetic precursors (Puffal et al., 2018). Nevertheless, there are many IMD-associated enzymes that are not involved in the cell envelope biosynthesis, suggesting more general functions of the IMD as a spatially distinct area of mycobacterial membrane, including the possible regulation of cytoplasmic metabolites, which is largely unexplored.

The biosynthetic enzymes for menaquinones (2-methyl-3-polyprenyl-1,4-naphthoquinones) are potential examples of such IMD-associated enzymes that are not directly involved in the cell envelope biosynthesis. Menaquinones are major lipoquinone electron carriers of mycobacterial respiratory chain. A major final product of the biosynthetic pathway is referred as MK-9 (II-H_2_), which carries a nonaprenyl chain with the second double bond (β-position) saturated (Collins et al., 1977). Its biosynthesis can be divided into the initial cytoplasmic reactions followed by the final membrane-associated steps (Meganathan, 2001). The membrane-associated reactions are mediated by three enzymes. First, the product of the cytoplasmic reactions, 1,4-dihydroxy-2-naphthoate, is attached to a polyprenol lipid by a membrane-bound polyprenyltransferase known as MenA (Dhiman et al., 2009) (Fig. 1A). Second, the resulting demethylmenaquinone is methylated on the aromatic ring by MenG (*syn.* MenH/UbiE), forming menaquinone (Dhiman et al., 2009). Finally, the double bond in the β-isoprene unit of the polyprenyl chain is reduced by the reductase MenJ to form the mature product, such as MK-9 (II-H_2_) (Upadhyay et al., 2015; 2018). Our comparative proteomic analysis of the IMD and the PM-CW suggested that MenG and MenJ are enriched in the IMD, while MenA was not detected in either the IMD or the PM-CW (Hayashi et al., 2016).

**Figure 1.**
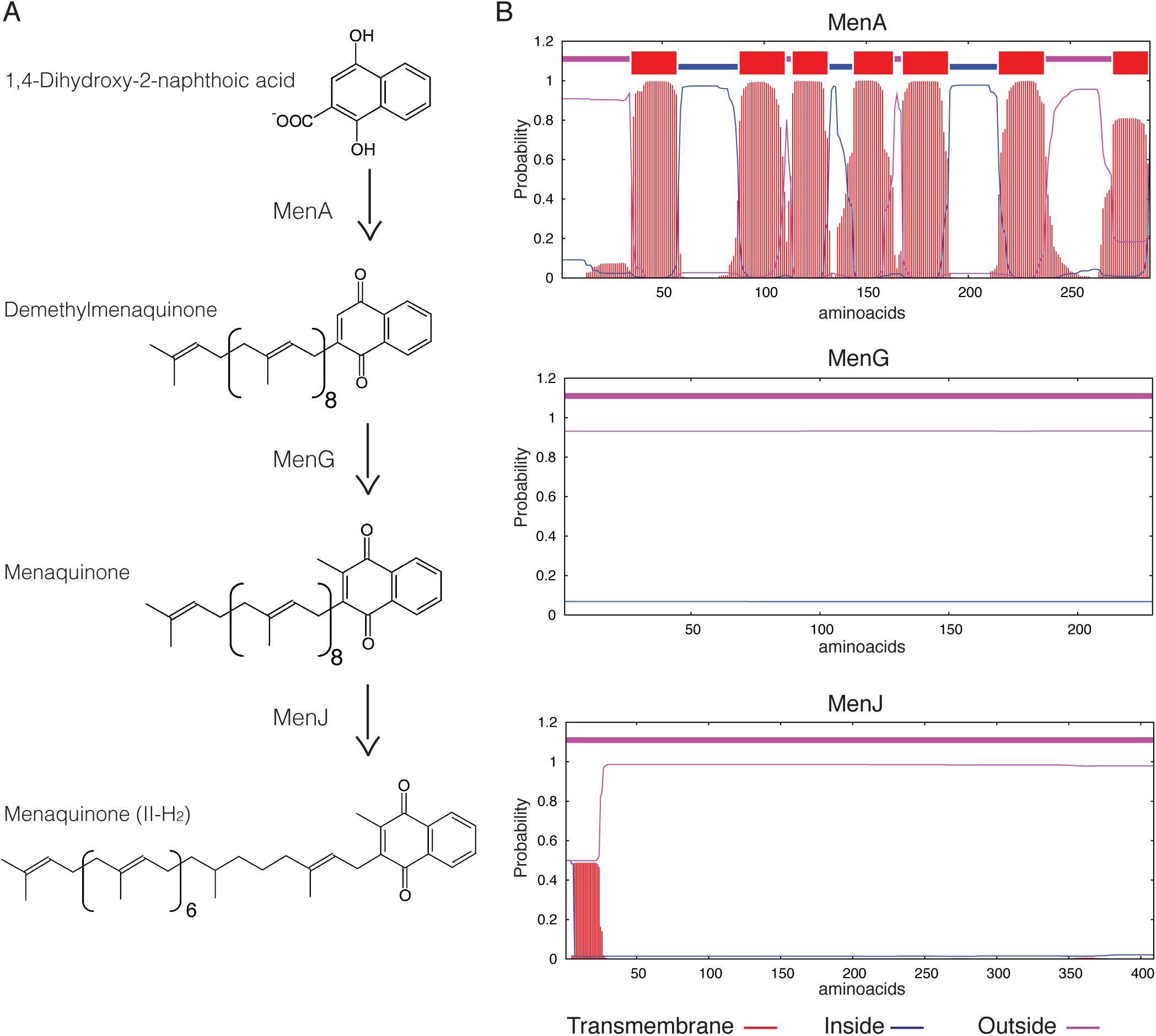
Last three steps of menaquinone biosynthesis in mycobacteria. (**A**) MenA adds a polyprenol such as nonaprenol to 1,4-dihydroxy-2-naphthoic acid forming DMK-9. MenG methylates the polar ring resulting in MK-9. MenJ reduces one C=C bond of the second prenyl group to form the mature MK-9 (II-H_2_). (**B**) Predicted transmembrane domains of the last menaquinone biosynthetic enzymes using TMHMM Server 2.0 (Krogh et al., 2001) based on amino acid sequence. MenA (upper panel) has seven predicted transmembrane helices, while MenG and MenJ show no predicted transmembrane domains.

Menaquinone biosynthesis is a critical process in mycobacteria. A previous study revealed Ro 48-8071 as an inhibitor of MenA, and demonstrated that this and other MenA inhibitors arrest the growth of both *Mycobacterium tuberculosis* and *M. smegmatis*, and reduce the cellular oxygen consumption (Dhiman et al., 2009). Another group showed that chemical inhibition of MenG is detrimental to the growth of *M. tuberculosis*, leading to the reduced oxygen consumption and ATP synthesis (Sukheja et al., 2017). In contrast, *menJ* is a dispensable gene in laboratory growth conditions: its deletion in *M. smegmatis* and *M*. *tuberculosis* produces viable mutants that show no significant changes in the growth rates (Upadhyay et al., 2015). Detailed analysis of this mutant revealed that the accumulation menaquinone-9 (MK-9) instead of MK-9 (II-H_2_) resulted in reduced electron transport efficiency. However, the mutant produced an increased amount of MK-9 to compensate partially for the loss of the mature species, indicating significant flexibility in meeting with the cellular needs of lipoquinones for respiration.

Combining evidence for the important roles of these enzymes with the new proteomic analysis suggesting that MenG and MenJ might be IMD-associated, we examined if the membrane steps of menaquinone biosynthesis is compartmentalized within the plasma membrane in *M. smegmatis*. In the present study, we directly demonstrated that MenG and MenJ are associated with the IMD while MenA is associated with the PM-CW. We further demonstrated that *menG* is an essential gene in *M. smegmatis*. Interestingly, partial depletion of MenG was detrimental to the *M. smegmatis* cells even though the cellular level of MK-9 and MK-9 (II-H_2_) remained high, implying a critical role of MenG in regulating menaquinone homeostasis in mycobacterial plasma membrane.

## Methods

### Cell cultures

*Mycobacterium smegmatis* mc^2^ 155 was grown as before (Hayashi et al., 2016) at 30°C in Middlebrook 7H9 broth supplemented with 11 mM glucose, 14.5 mM NaCl, and 0.05% Tween-80, or at 37°C on Middlebrook 7H10 agar supplemented with 11 mM glucose and 14.5 mM NaCl. When required, the medium was supplemented with 100 µg/ml hygromycin B (Wako), 50 µg/ml streptomycin sulfate (Fisher Scientific), 20 µg/ml kanamycin sulfate (MP Biochemicals), or 5% sucrose.

### Construction of plasmids

Plasmids used in this study are summarized in Table S1.

pMUM040 – To create expression vector for MenA which is C-terminally tagged with a hemagglutinin (HA) epitope, the gene was amplified by PCR (Table S2) using primers carrying appropriate restriction enzyme sites. The product was digested with BspEI and ligated to the vector backbone of pMUM038, which was linearized by BspEI/SspI double-digestion. The pMUM038 vector is identical to pMUM011 (Hayashi et al., 2016), a derivative of pVV16, but its sole NdeI site was removed by linearizing the plasmid using NdeI, blunting using the T4 polymerase, and circularizing using a DNA ligase.

pMUM042 – To create expression vector for MenG, which is C-terminally tagged with an HA epitope, the PCR product (Table S2) was inserted by blunt-end ligation to the vector backbone of pMUM012 (Hayashi et al., 2016), linearized by EcoRV and ScaI.

This intermediate plasmid, pMUM039, was then double-digested with NdeI and ScaI, and the fragment carrying *menG* gene was ligated into the linearized vector backbone of pMUM040 digested with the same enzymes.

pMUM055 – To knockout the endogenous *menG* gene, we amplified upstream and downstream regions of *menG* using the primers shown in Table S2 and digested with Van91I and DraIII, respectively. The two fragments were then ligated into Van91I-digested pCOM1 as previously described (Hayashi et al., 2016). The resulting plasmid, pMUM055, was used for allelic exchange of *menG* in *M. smegmatis* via a two-step recombination process as previously described (Hayashi et al., 2016; Rahlwes et al., 2017). The deletion of the *menG* gene was confirmed by PCR using primers A312 and A313 (Table S2).

pMUM098 – To create a MenG-HA expression vector with a kanamycin resistance selection marker, *menG-HA* gene fragment was isolated from pMUM042 (see above) by XmnI/EcoRI double digestion, and was inserted to XmnI/EcoRI double-digested pMUM087. pMUM087 is an NdeI-free version of pMV361 (Stover et al., 1991) (gift from Dr. William R. Jacobs Jr., Albert Einstein College of Medicine), created by digesting pMV361 with NdeI, and blunt-ending and re-ligating the linearized fragment.

pMUM103 – To create an expression vector for MenJ-HA, the gene was amplified by PCR (Table S2), and the PCR product was inserted directionally to pMUM098, from which the preexisting insert was removed by NdeI/ScaI double digestion.

pMUM058 – To create an expression vector for MenG tagged with mTurquoise, the mTurquoise gene was amplified by PCR (Table S2) from pYAB281 containing mTurquoise (Hayashi et al., 2016). The PCR product was then digested with ScaI and inserted to pMUM042, which was linearized by the same enzyme, creating an expression vector for C-terminally mTurquise-HA epitope-tagged fusion protein.

pMUM119 – To create a dual-control tet-off expression vector for MenG, in which the protein is fused with HA epitope and DAS degradation tag at the C-terminus, we first created an intermediate construct pMUM106 by Gibson assembly of ClaI/NdeI double-digested pDE43-MCS (Blumenthal et al., 2010), the PCR amplified promoter region of pEN12A-P766-8G (A442/A443, Table S2) (Kim et al., 2013) (gift from Dr. Christopher Sassetti, University of Massachusetts Medical School), and the PCR-amplified *menG* gene from pMUM090 (A444/A445, Table S2), resulting in a MenG expression vector driven by the weak P766-8G promoter. We then inserted a fragment for the expression of TetR38, which was PCR-amplified from pEN41A-T38S38 (A487/A488, Table S2) (Kim et al., 2013) and digested with EcoRV and BspTI, into pMUM106 digested with EcoRV and BspTI, resulting in pMUM110. To attach the HA and DAS tags, the *menG-HA* fragment was amplified by PCR (A185/A506, Table S2) from pMUM098. The PCR fragment and pMUM110 were digested with SacI and VspI and ligated, creating pMUM119. The SspB expression vector (pGMCT-3q-taq25) and non-replicative integrase expression vector (pGA-OX15-int-tw) (gift from Dr. Christopher Sassetti, University of Massachusetts Medical School) were co-electroporated to allow stable integration of pGMCT-3q-taq25.

Plasmid constructs (Table S1) were electroporated into *M. smegmatis* for integration and homologous recombination as previously described (Hayashi et al., 2016).

### Density gradient fractionation and protein analysis

Log phase cells (OD_600_ = 0.5-1.0) were pelleted, lysed by nitrogen cavitation, and subjected to sucrose density fractionation as previously described (Hayashi et al., 2016). Briefly, 1.2 ml of the lysate was loaded on top of a 20-50 % sucrose gradient prepared in a 14 x 95 mm tube (Seton Scientific). The gradient was spun at 35,000 rpm (218,000 x *g*) for 6 h at 4°C in a SW-40 rotor (Beckman-Coulter). Thirteen 1-ml fractions were then collected and used for further biochemical analysis. Protein concentration was determined by the bicinchoninic acid (BCA) assay (Pierce). Sucrose density was determined by a refractometer (ATAGO). For SDS-PAGE and western blotting, an equal volume of each fraction was loaded as described before (Hayashi et al., 2016; 2018).

### Fluorescence microscopy

Fluorescence microscopic live imaging was done as previously described (Hayashi et al., 2016).

### Plasmid swap

Expression vectors, pMUM098 (*men G-HA*, Kan^r^) and pMUM087 (empty vector, Kan^r^), were electroporated into *M. smegmatis* Δ*menG* L5::*men G-HA* Str^r^ strain to swap the inserted plasmid at the L5 integration site. The swapping was verified by culturing the transformed colonies on Middlebrook 7H10 plates containing kanamycin or streptomycin.

### *menG* conditional knockdown

*M. smegmatis* Δ*menG* L5::*men G-HA* Kan^r^ was transformed with the plasmid pMUM119 (*tet*_*Off*_ *men G-HA-DAS* Str^r^) to swap at the L5 integration site to create Δ*menG L5*::*tet*_*Off*_ *men G-HA-DAS* Str^r^. This new strain was then transformed with pGMCT-3q-taq25/pGA-OX15-int-tw (*tet*_*On*_ *sspB* Kan^r^), an integrative plasmid that recombines at an *attB* site for the mycobacteriophage Tweety (Pham et al., 2007), resulting in the *menG* dual-switch knockdown strain, Δ*menG L5*::*tet*_*Off*_ *men G-HA-DAS* Str^r^ *Tweety*::*tet*_*On*_ *sspB* Kan^r^.

A starter culture of the dual-switch *menG* knockdown strain, grown in Middlebrook 7H9 medium containing streptomycin and kanamycin, was inoculated into fresh Middlebrook 7H9 medium with or without 100 ng/ml anhydrotetracycline (ATC) and subsequently sub-cultured by 100-fold dilution every 24 h. One ml of each culture was taken for OD_600_ reading to monitor the growth, and colony formation unit (cfu) was determined at the 72-h timepoint. As controls, we used *M. smegmatis* strains carrying *Tweety*::*tet*_*On*_ *sspB* Kan^r^ alone. For menaquinone-4 (MK-4) supplementation, we prepared 80 mM MK-4 stock solution in dimethyl sulfoxide and slowly added to a culture to achieve a final concentration of 400 µM, following a previously published protocol (Dhiman et al., 2009). To analyze the protein depletion kinetics, western blot images were recorded and quantified using ImageQuant LAS4000mini (GE Healthcare).

### Mass spectrometric analysis of lipids

A previously reported comparative lipidomics dataset (reproduced in Fig. S2A) (Hayashi et al., 2016) was further analyzed for annotations of several menaquinone species based on mass, which were subjected to validation by collision-induced dissociation mass spectrometry. For targeted analysis of menaquinone, we grew the cells in the presence and absence of ATC for 72 h, sub-culturing at every 24 h and harvested cells at 72 h. Cells were lysed by nitrogen cavitation and the lipids were extracted from the whole cell lysate. For lipid extraction, 500 µl of cell lysates were supplemented with 10 nmol of MK-4 (Millipore-Sigma) as an internal standard. Six ml ice-cold 0.2 M perchloric acid in methanol was added along with 6 ml petroleum ether (preheated to 40-60°C) as described previously (Bekker et al., 2007). The mixture was vortexed and spun, and the top organic layer was transferred to a new tube. The lower layer was extracted again with 6 ml of petroleum ether and the organic extracts were combined. The combined organic extract was washed once with 6 ml of water, and the final top organic layer was transferred to a new tube, dried and resuspended in 100 µl chloroform/methanol (1:1). To evaluate the extraction efficiency, we subjected 10 µl to thin layer chromatography and orcinol/H_2_SO_4_ staining for the detection of phosphatidylinositol mannosides (Morita et al., 2004). AcPIM2 bands were quantified using Fiji (Schindelin et al., 2012), and used to adjust the lipid concentration of each sample.

The purified lipids were subjected to high-performance liquid chromatography (HPLC)-tandem mass spectrometry (Orbitrap Fusion with higher energy collisional dissociation (HCD) coupled with UltiMate 3000 HPLC system, Thermo Scientific), using PC-HILIC column (Shiseido) with acetonitrile/water (95:5) with 10 mM ammonium acetate (pH 8.0) as the mobile phase. The ESI was operated in a positive polarity mode, with spray voltage of 2.8 kV and flow rate of 0.3 ml/min. The full scan range was 100 to 1,200 *m/z* and the data was recorded using Xcalibur 3.0.63 software package (Thermo Scientific). For HCD, a quadrupole isolation mode was used with collision energy of 40±5% and data detected by Orbitrap (Thermo Scientific). The targeted *m/z* were defined as 771.6075 for demethylmenaquinone-9 (DMK-9), 785.6231 for MK-9, 787.6388 for MK-9 (II-H_2_), and 445.3101 for MK-4. The detection efficiency of MK-9 relative to MK-4 was determined using 20 pmol of commercially available MK-9 (Santa Cruz biotechnology) and MK-4 (Millipore-Sigma).

### Oxygen consumption

The effect of MenG depletion on oxygen consumption was evaluated by methylene blue decolorization. One OD unit of each culture from the 72-h time-point was harvested, resuspended in 2 ml of Middlebrook 7H9, and supplemented with 0.001% methylene blue (Ricca). In sealed cuvettes, oxygen consumption was monitored by absorbance at 665 nm.

### Measurement of cellular ATP levels

The dual-switch *menG* knockdown strain was incubated in the presence and absence of ATC for 72 h, sub-culturing at every 24 h, as described above. Intracellular ATP was determined by BacTiter Glo microbial cell viability assay (Promega), following manufacturer’s instruction.

## Results

### The maturation of MK-9 takes place in the IMD

The three final steps on MK-9 biosynthesis are catalyzed by the enzymes MenA, MenG and MenJ (Fig. 1A). MenA is a protein with multiple predicted membrane spanning domains (Fig. 1B). MenG and MenJ have no predicted transmembrane domains, so the patterns and specific mechanisms of membrane association of these proteins might vary. Previously, we showed by comparative proteomics that peptide fragments corresponding to known MenG and MenJ sequences were recovered at higher level in the IMD than in the conventional plasma membrane (PM-CW) (Hayashi et al., 2016). Thus, MenG and MenJ are potentially IMD-associated proteins peripherally bound to the membrane surface. However, our proteomic analysis did not examine the cytoplasmic fraction, and therefore cannot exclude the possibility that these proteins reside also in the cytoplasm.

To determine the subcellular localization of these three enzymes based on direct detection of intact proteins in all three compartments, we expressed MenA-HA (expected molecular weight, 29 kDa), MenG-HA (25 kDa) and MenJ-HA (44 kDa) individually at the site-specific integration site of mycobacteriophage L5 in *M. smegmatis*, and confirmed that all three proteins were expressed at the expected molecular weight (Fig. S1). We then performed sucrose density gradient fractionation of each strain, and determined the subcellular localization of these enzymes within the gradient. PimB’ and MptA are the protein markers for the IMD and PM-CW, respectively, and MenA-HA was enriched in the fractions corresponding to the PM-CW together with MptA (Fig. 2A). In contrast, MenG and MenJ were enriched in the IMD (Fig. 2B-C). The low density fractions (Fr. 1-2), that are high in total protein content, are known to be enriched in cytoplasmic proteins (Hayashi et al., 2016; Morita et al., 2005). Neither MenG nor MenJ was found in the cytoplasmic fraction, indicating that these two proteins are stably associated with the IMD.

**Figure 2.**
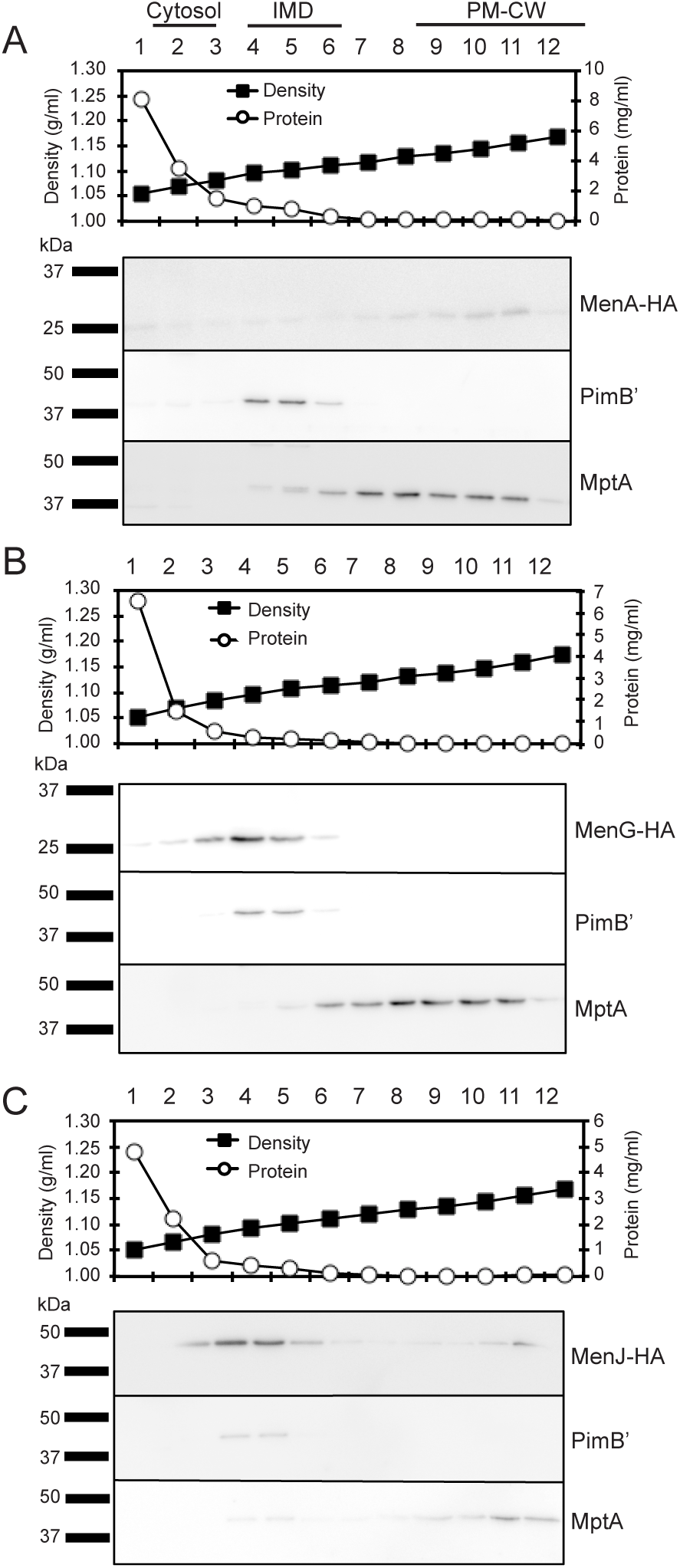
Subcellular localization of menaquinone biosynthesis. (**A-C**) Sucrose density gradient fractionation of cell lysates prepared from strains expressing (**A**) MenA-HA, (**B**) MenG-HA and (**C**) MenJ-HA. Protein concentration and sucrose density in each fraction were plotted in the graph. Protein markers for the IMD and the PM-CW were PimB’ (41 kDa) and MptA (54 kDa), respectively. All experiments were done more than twice and representative data are shown.

To determine the subcellular localization of MenG in live bacteria, we next introduced an MenG-mTurquoise-HA expression vector, and expressed the fluorescent fusion protein in a previously established *M. smegmatis* strain expressing HA-mCherry-GlfT2 from the endogenous *glfT2* locus. GlfT2 is a galactosyltransferase involved in the arabinogalactan precursor synthesis, and is an IMD-associated protein (Hayashi et al., 2016). We first confirmed by sucrose density gradient that MenG-mTurquoise-HA co-fractionates with HA-mCherry-GlfT2 and PimB’ (Fig. 3A), but not with the PM-CW marker, MptA. The expected molecular weight of HA-mCherry-GlfT2 and MenG-mTurquoise-HA are 100 and 50 kDa, respectively, allowing separate detection of these two HA-tagged proteins in a single western blot. This result also revealed the relatively lower expression level of MenG-mTurquoise-HA in comparison to HA-mCherry-GlfT2, even though the expression of MenG is driven by a strong promoter. Consistent with the apparently lower MenG expression in cellular extracts, we also observed a much weaker level of fluorescence from mTurquoise by fluorescence microscopy live imaging (Fig. 3B). Although the weak fluorescence and a higher background due to autofluorescence (Patiño et al., 2008)(Fig. 3C) made the image analysis difficult, we were able to observe the polar enrichment of MenG-mTurquoise-HA, which correlated with the polar enrichment of HA-mCherry-GlfT2 (Fig. 3B).

**Figure 3.**
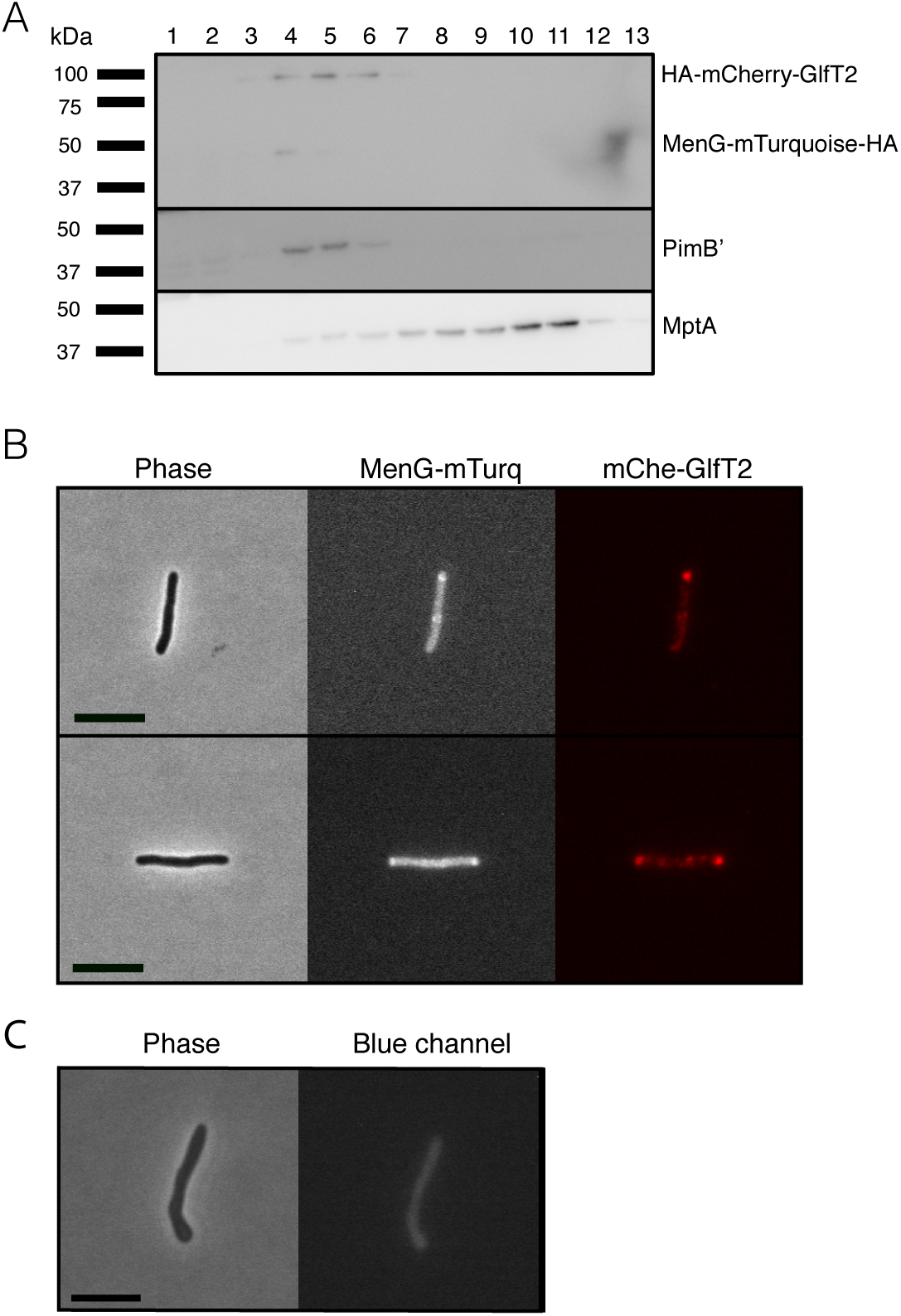
Co-localization of MenG with IMD associated protein GlfT2. (**A**) Sucrose density fractionation of strain expressing HA-mCherry-GlfT2 (100 kDa) and MenG-mTurquoise-HA (50 kDa). The epitope-tagged proteins were detected by anti-HA antibody. PimB’ (41 kDa) and MptA (54 kDa), respectively, indicate the IMD and PM-CW fractions. (**B**) Fluorescence microscopy showing localization of both MenG-mTurquoise-HA and HA-mCherry-GlfT2 at the pole of growing *M. smegmatis* cells. (**C**) Autofluorescence of WT *M. smegmatis* on blue channel observed under the identical image acquisition setting as in panel B. Scale bar = 5 µm. All experiments were done more than twice and representative data are shown.

To determine if any menaquinone species are enriched in the IMD, we analyzed a comparative HPLC time-of-flight (TOF) mass spectrometry-derived lipidomic dataset comprised of 11,079 separately detected molecular events (Hayashi et al., 2016). This method was previously validated to extract hydrophobic molecules, including menaquinones, and normal phase chromatography reduces cross-suppression by more polar species, allowing semiquantitative detection of lipid compounds (Lahiri et al., 2016; Layre et al., 2011). The IMD preparations were previously validated based on IMD-specific proteins and revealed IMD-associated phospholipids among compounds upregulated in the IMD. The majority of these compounds were unnamed, but are discoverable based on matching their *m/z* values with the MycoMap dataset (Layre et al., 2011). This approach allowed the identification of signals matching the *m/z* value of DMK-9 and MK-9 in their reduced and non-reduced forms. While DMK-9 was equally present in both sites, the MK-9 species were overexpressed in IMD to varying degrees (Fig. S2A-B). The identity of both MK-9 (II-H_2_) and MK-9 were confirmed by collision-induced dissociation mass spectrometry (Fig. S2C), showing that they are in the ketone form with a reduced double bond in the nonaprenyl lipid moiety of MK-9 (II-H_2_). Taken together, these data suggest that 1) MenA produces DMK-9 in the PM-CW; 2) DMK-9 relocates from the PM-CW to the IMD; 3) MenG methylates DMK-9 to generate MK-9 in the IMD; and 4) MenJ reduces the prenyl lipid of MK-9 to form the mature molecule, MK-9 (II-H_2_), in the IMD (Fig. 4). For MK-9 (II-H_2_) to function as an electron carrier, it may then have to relocate back to the PM-CW because the respiratory chain enzymes are found in the PM-CW (Hayashi et al., 2016).

**Figure 4.**
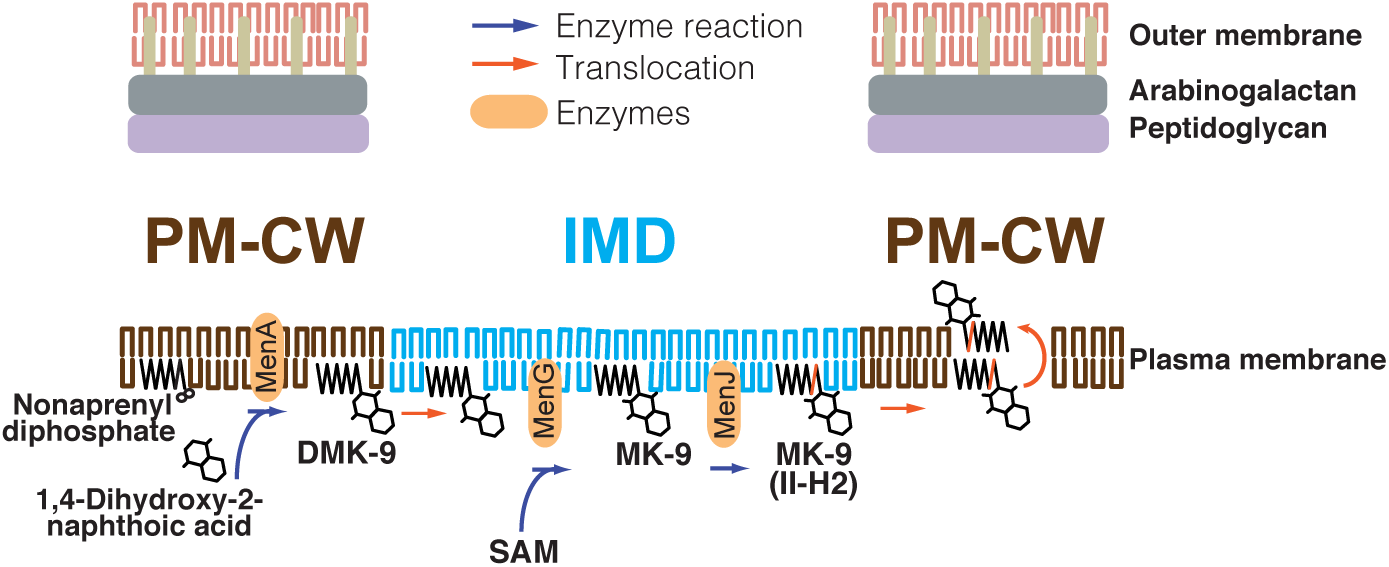
Proposed spatial compartmentalization of MK-9 biosynthetic pathway. DMK-9 is formed in the PM-CW by the prenyltransferase MenA. DMK-9 traffics to the IMD and modified by MenG and MenJ, forming MK-9 and MK-9 (II-H_2_), respectively. The mature molecule can then be transferred to the PM-CW to serve as an electron carrier. The red line in the polyprenol moiety of MK-9 (II-H_2_) indicates the saturation of the second isoprene unit mediated by MenJ. SAM, S-adenosylmethionine.

### The *menG* gene is essential in *M. smegmatis*

The intricate spatial segregation of biosynthetic enzymes suggests that menaquinone biosynthesis may be a highly regulated process. The association of this pathway also implies an indirect role of the IMD in the central energy metabolism. Nevertheless, little is known why the MenG- and MenJ-dependent modifications on DMK-9 are physiologically important. MenJ is dispensable for growth of both *M. tuberculosis* and *M. smegmatis* in standard laboratory growth conditions (Upadhyay et al., 2015). In contrast, *menG* is predicted to be essential in *M. tuberculosis* (Griffin et al., 2011), but no direct or indirect information about its essentiality is available for *M. smegmatis*.

To begin delineating the function of MenG, we first attempted to knock out *menG* by a markerless deletion using a plasmid that carries *sacB* gene as a negative selection marker and hygromycin resistance gene as a positive selection marker (Hayashi et al., 2016) (Fig. S3A). We confirmed the establishment of a single-crossover mutant that is sensitive to sucrose (due to *sacB* gene) and resistant to hygromycin. We then grew the single-crossover mutant in nonselective medium to allow the second crossover event, and isolated 17 colonies that are resistant to sucrose and sensitive to hygromycin. When we analyzed these double-crossover candidates, they were all found to be wild-type revertants and no candidate had the *menG* deletion (data not shown). These initial observations suggested that *menG* is an essential gene.

To test this further, we created a merodiploid strain of the single-crossover mutant, in which a *menG* expression vector with a streptomycin resistance marker (pMUM042) was inserted at the L5 integration site (Fig. S3A). We successfully isolated double-crossover mutants from the merodiploid single-crossover strain, as confirmed by PCR of the endogenous *menG* gene locus (Fig. S3B). Using the double-crossover mutant, we attempted to swap the *menG* expression vector, pMUM042, carrying streptomycin resistance marker with another L5-integrative *menG* expression vector, pMUM098, carrying kanamycin resistance marker or with an empty vector, pMUM087, carrying kanamycin resistance marker as a control. When pMUM098 was used, 240 colonies were obtained (Fig. S4). We patched 78 colonies on Middlebrook 7H10 medium containing either streptomycin or kanamycin, and found that all 78 colonies were sensitive to streptomycin and resistant to kanamycin, suggesting that pMUM042 was swapped with pMUM098. In contrast, when the empty vector pMUM087 was used, only 3 colonies were obtained and they were all resistant to both kanamycin and streptomycin, suggesting that the cells could not lose pMUM042 carrying *menG* gene. Using a newly established kanamycin-resistant strain carrying pMUM098, we attempted to swap back using streptomycin-resistant pMUM042. Again, we were able to isolate 20 colonies using pMUM042, but no legitimate swapping occurred using the empty vector, pMUM038 (Fig. S4). Taken together, these data strongly support that *menG* is an essential gene.

### MenG depletion leads to growth arrest without significant depletion of menaquinone

To examine the mechanistic basis of MenG essentiality in *M. smegmatis*, we constructed a cell line with a dual-control switch in which ATC suppresses the expression of *menG* and degrades MenG protein simultaneously (Fig. 5A) (Kim et al., 2013). In this Δ*menG L5*::*tet*_*Off*_ *men G-HA-DAS* Str^r^ *Tweety*::*tet*_*On*_ *sspB* Kan^r^ strain, an inducer ATC turns off the transcription of *men G-HA-DAS* gene. At the same time, the transcription of *sspB* gene is turned on, and the SspB adaptor protein recognizes and targets the DAS-tagged protein for ClpXP-dependent degradation. Upon addition of ATC, the cells started to show deficiency in growth after two consecutive series of 24-h sub-culturing. The OD_600_ reading for the treated cells became significantly lower after 3 rounds of sub-culturing (Fig. 5B). The cfu for untreated and ATC-treated cultures were 1.8 x 10^7^ and 5.2 x 10^6^ cfu/ml, respectively, comparable to the OD measurements, suggesting MenG depletion is bacteriostatic rather than bactericidal. We examined the protein level of MenG over the time course, and found that total protein fell to ~76% of the level found in untreated cells by 48 h (Fig. 5C). This moderate suppression of MenG continued even at the 72 h time point, where the MenG protein was reduced further to the ~35% of the level found in untreated cells (Fig. 5C). The relatively mild MenG depletion made us wonder if the lack of growth is due to the depletion of menaquinone. In *M. tuberculosis*, the growth arrest by the MenG inhibitor was rescued by the addition of MK-4 as a surrogate menaquinone (Sukheja et al., 2017). Therefore, we added MK-4 to see if exogenously added menaquinone can rescue the growth of the mutant in the presence of ATC. As shown in Fig. S5, MK-4 supplementation was unable to rescue the growth of the ATC-treated cells, suggesting that the growth defect of the mutant might not be due to the depletion of menaquinone.

**Figure 5.**
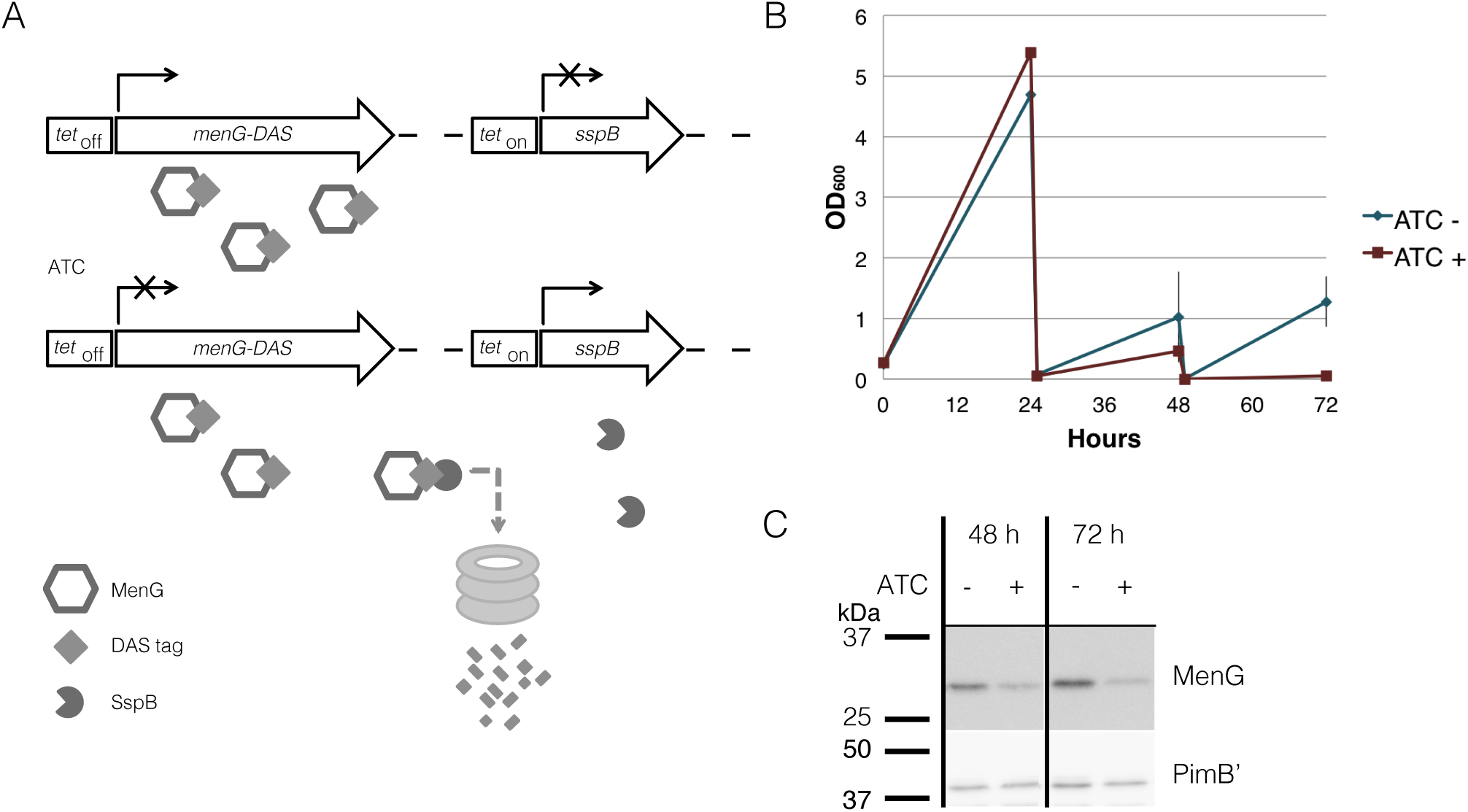
MenG knockdown. (**A**) Scheme of MenG depletion. When the Δ*menG L5*::*tet*_*Off*_ *men G-HA-DAS* Str^r^ *Tweety*::*tet*_*On*_ *sspB* Kan^r^ strain is exposed to ATC, a tetracycline analog, MenG expression is shut off and the protein is tagged for degradation by SspB. (**B**) Growth curve of Δ*menG L5*::*tet*_*Off*_ *men G-HA-DAS* Str^r^ *Tweety*::*tet*_*On*_ *sspB* Kan^r^ exposed to ATC over 72 hours of subculturing every 24 hours. The averages of biological triplicates are shown with standard deviations. (**C**) MenG depletion after 48 and 72 hours of ATC treatment detected by western blotting. Images were captured by the ImageQuant LAS4000mini image documentation system and bands were quantified using ImageQuant analysis software (GE Healthcare). ATC, anhydrotetracycline. All experiments were done more than twice and representative data are shown.

To evaluate the impact of MenG depletion on cellular menaquinone levels, we took the 72-h time point, and performed HPLC tandem mass spectrometry analysis on the lipid extracts from crude lysates. We confirmed the identity of MK-9 (*m/z* 785.6231), MK-9 (II-H_2_) (787.6388), and DMK-9 (771.6075) by fragmentation (Fig. S6), and quantified the levels of each species relative to the internal standard MK-4 (*m/z* 445.3101).

As expected, we saw a significant increase in the DMK-9 levels when cells were treated with ATC (Fig. 6A). Surprisingly, the levels of MK-9 and MK-9 (II-H_2_) were not significantly different between the untreated and MenG-depleted strains (Fig. 6B-C). These data support the idea that the partial depletion of MenG leads to the accumulation of the MenG substrate, DMK-9, immediately impacting cellular metabolic activities prior to affecting the cellular levels of MK-9 and MK-9 (II-H_2_).

**Figure 6.**
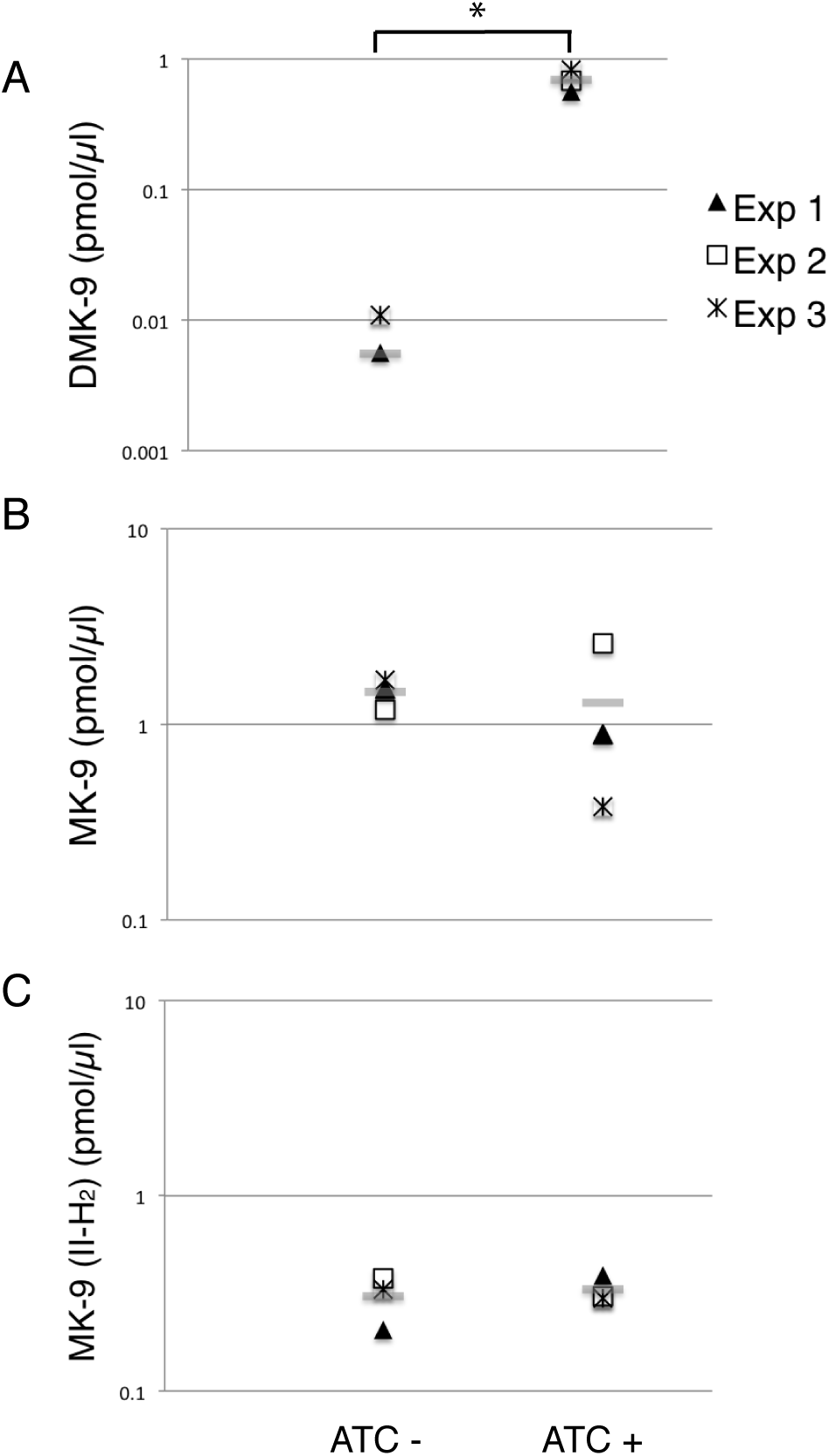
Changes in menaquinone species upon MenG depletion. Lipid extracts from crude cell lysates were analyzed by HPLC mass spectrometry to quantify (**A**) DMK-9, (**B**) MK-9 and (**C**) MK-9 (II-H_2_). In three independent experiments, lysates were prepared after 72-h growth with or without ATC (biological triplicates). From each replicate of the biological triplicates, lipids were extracted and analyzed twice (technical duplicates). MK-4 was added as an internal standard to control the efficiency of lipid extraction and HPLC mass spectrometry analysis. Each point in the graphs is the average of the technical duplicate, and the grey line represents the average of biological triplicates. The unit is pmol of indicated menaquinone species per µl of cell lysate. *, p < 0.05 by t-test.

### Impact of MenG depletion on respiration and cellular ATP levels

Because MenG depletion appears to have no immediate effect on the levels of MK-9 and MK-9 (II-H_2_), we examined if cellular respiration is affected upon MenG depletion. Cells were grown as previously for 72 h with sub-culturing at every 24 h, and aliquots of cell suspension was tested for O_2_ consumption using the decolorization of methylene blue. We found that the untreated (ATC-) cells rapidly depleted the O_2_ from the media, but the treated (ATC+) cells consumed very little O_2_ during the same period (Fig. 7A), suggesting that respiration in the ATC+ cells is significantly reduced even though these cells are viable as indicated above by the cfu.

**Figure 7.**
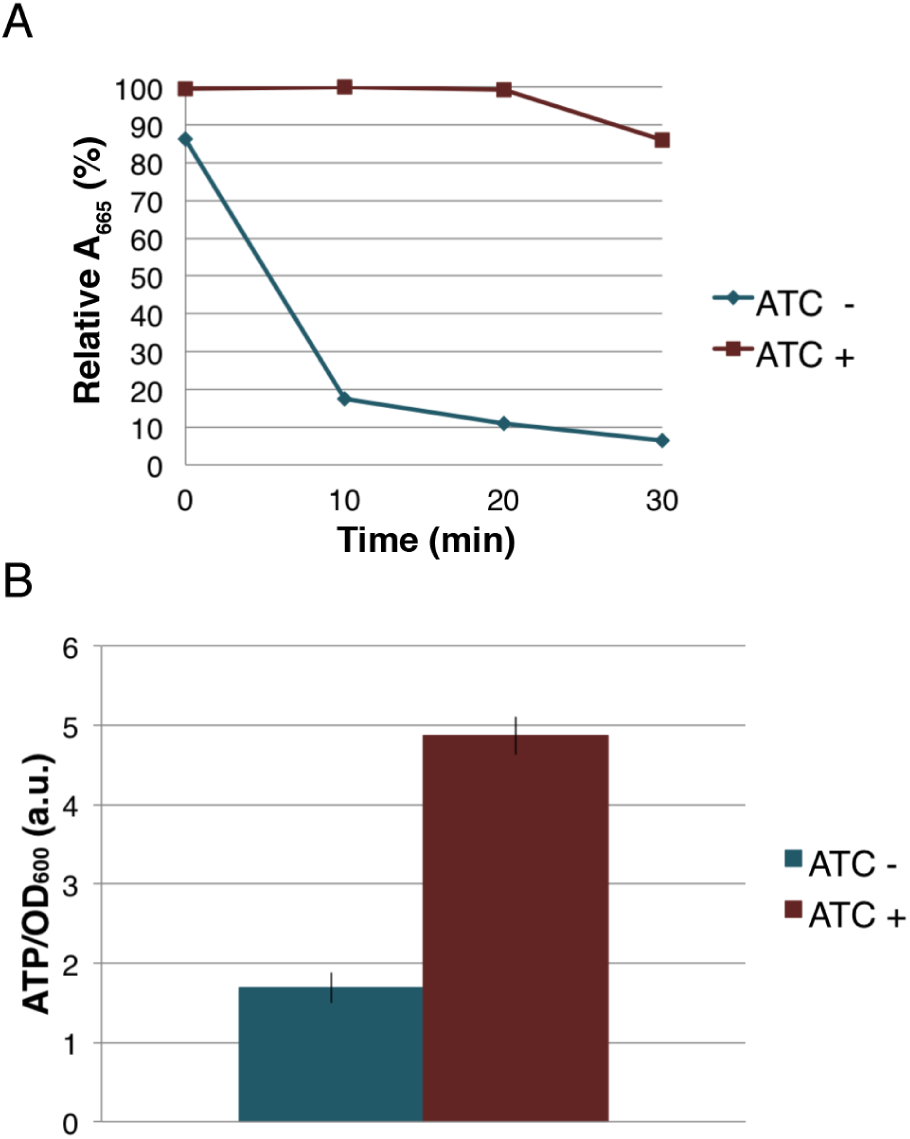
Effect of MenG depletion on aerobic respiration and intracellular ATP level. (**A**) Oxygen consumption by *M. smegmatis* MenG depletion strain after 72-h with and without treatment with ATC. The decolorization of methylene blue in the media was used as an indication of the oxygen consumption, taking the A_665_ of methylene blue immediately after the addition of ATC (at 0 min) as 100%. (**B**) ATP accumulation over a 24-hour period during the third sub-culturing from the 48-h to the 72-h time point with and without ATC. The averages of biological triplicates are shown with standard deviations. ATC, anhydrotetracycline.

The severe reduction in the rate of aerobic respiration suggested an impact on the central metabolism. We next examined the cellular level of ATP when the cells were treated with ATC during the third sub-culturing from the 48-h time point to the 72-h time point. We found that the MenG-depleted (ATC+) cells accumulated ~3 times more ATP that the untreated (ATC-) cells (Fig. 7B). This was not due to the SspB expression because a control cell line, which only expresses SspB upon ATC addition, did not show any changes in cellular ATP levels (Fig. S7). These data are consistent with the idea that MenG depletion resulted in the reduction of the cellular metabolism, and the lack of energy consumption resulted in the accumulation of ATP.

## Discussion

The IMD is a metabolically active membrane domain that mediates many distinct biosynthetic pathways. In this study, we demonstrated that the final maturation steps of the menaquinone biosynthesis take place in the IMD, and MenG, one of the IMD-associated enzymes, is essential for the growth of *M. smegmatis*. The IMD association of MenG is supported by three lines of evidence gathered *in vitro* and in live cells. First, proteomic analysis indicated that MenG is more enriched in the IMD than in the PM-CW (Hayashi et al. 2016). Second, epitope-tagged MenG was biochemically localized to the IMD by density gradient fractionation. Although we cannot completely rule out the possibility that the HA epitope tag interferes with the subcellular localization, the IMD localization of the fusion protein was consistent with the proteomic identification of the endogenous protein in the IMD as mentioned above. Finally, fluorescent protein-tagged MenG showed colocalization with a known IMD marker at polar regions of actively growing cells by fluorescence microscopy. Combined with the IMD localization of epitope-tagged MenJ, we suggest that menaquinone species, MK-9 and MK-9 (II-H_2_), are produced in the IMD. Indeed, the comparative lipidomic analysis suggested that MK-9 (II-H_2_) as well as MK-9 are relatively enriched in the IMD, but overall do not show the high levels of segregation as seen for the proteins that act on them. These observations suggest cellular regulation of the enzymes with substrates diffusing between both sites.

Do menaquinones have a functional role in the IMD or is it merely produced there? We propose that menaquinones function as an electron carrier for some IMD-associated enzymes. For example, we have previously shown that the dihydroorotate dehydrogenase PyrD, an enzyme involved in pyrimidine biosynthesis, is an IMD-associated protein (Hayashi et al., 2016). Mycobacterial PyrD is a member of the class 2 dihydroorotate dehydrogenases (Björnberg et al., 1997; Munier-Lehmann et al., 2013), which utilize quinones instead of NADH as an electron acceptor. Therefore, *de novo* synthesized menaquinones are locally available to support the IMD-resident PyrD reaction.

Nevertheless, a major fraction of menaquinones must also be available for cytochromes in the respiratory chain. Our comparative proteomic analysis suggested that the respiratory chain cytochromes as well as H^+^-ATPases are enriched in the conventional plasma membrane (Hayashi et al., 2016). For example, the subunits of cytochrome *c* reductase (QcrCAB; MSMEG_4261-4263) and aa_3_ cytochrome *c* oxidase (CtaC; MSMEG_4268), as well as the subunits of H^+^-ATPases (*e.g.* alfa, beta, H, F and A; MSMEG_4938, MSMEG_4936, MSMEG_4939, MSMEG_4940, MSMEG_4942, respectively) are enriched in the PM-CW proteome. Furthermore, the major NADH oxidase reactions take place in the PM-CW (Morita et al., 2005). Together, menaquinones produced in the IMD may be relocated to the PM-CW to support cellular respiration. Whether menaquinones diffuse through different membrane areas or require a transport mechanism remains an important question to be addressed in the future.

Several independent lines of experimental evidence clearly indicated that MenG is an essential protein in *M. smegmatis*. In the dual-switch knockdown system, the depletion of MenG protein was only partial even after three consecutive 24-h subcultures. We do not know why this mutant shows this unusual protein depletion kinetics, but speculate that the protein degradation is not efficient and MenG might have a prolonged half-life. Nevertheless, this mild MenG depletion led to the growth arrest. Why is this mild MenG depletion detrimental to *M. smegmatis*? Indeed, MK-9 is still abundantly present after three 24-h subcultures with ATC induction. The MenG substrate, DMK-9, however, showed a significantly increase in the treated population. We speculate that MenG might play a key regulatory role in the IMD, and the disruption of the balance between MK-9 and DMK-9 by its partial depletion could induce metabolic shutdown and the cessation of growth.

In *M. tuberculosis*, a recent study demonstrated that MenG is an effective drug target, and its inhibition led to the reduced oxygen consumption and ATP production. Our data in *M. smegmatis* is consistent with the previous findings in *M. tuberculosis* in that MenG is an essential protein, but also illuminate some important differences. First, we could not rescue the MenG depletion by the addition of MK-4, while exogenously supplemented MK-4 was apparently incorporated into the plasma membrane to function as a surrogate electron carrier in *M. tuberculosis* in the presence of MenG inhibitor (Sukheja et al., 2017). Second, MenG depletion in *M. smegmatis* did not lead to the reduction in ATP production. These differences could possibly be attributed to the differing chemical versus genetic methods of perturbation used in the two studies. Importantly, DMK, which accumulates upon MenG perturbation, is a fully functional electron carrier in *Escherichia coli* (Sharma et al., 2012; Unden and Bongaerts, 1997; van Beilen and Hellingwerf, 2016), implying that the physiological importance of the MenG-mediated methylation of the DMK aromatic ring in mycobacteria is not merely a matter of the mid-point electron potential of quinones as electron carriers.

Why does ATP accumulate during MenG depletion? In many other bacteria, when proton gradient formation is compromised, ATP synthase can be reversed to hydrolyze ATP and used to reestablish the proton gradient (Ballmoos et al., 2009). In mycobacteria, on the other hand, such reverse action of ATP synthetase is blocked and cannot be used to energize the membrane (Haagsma et al., 2010). Therefore, even when the cells are exposed to hypoxic conditions and cannot create a sufficient level of proton motive force, the accumulating ATP in the cell might not be utilized for energizing the membrane.

MenG expression is upregulated in response to the depletion of S-adenosylmethionine, indicating one example of transcriptional regulations of *menG* gene in response to changing metabolic state of the cell (Berney et al., 2015). We speculate that MenG depletion might be mimicking an adaptive response to an environmental change, leading the cells to stop aerobic respiration and consumption of ATP. In addition, we cannot rule out the possibility that the cells start using an alternative electron acceptor instead of oxygen. Such an adaptive response is known in *E*. *coli* (Edwards et al., 2006; Georgellis et al., 2001; Malpica et al., 2004), where changes in the environmental oxygen level result in a switch of lipoquinone species used in the electron transport chain. In this regard, when oxygen is depleted in mycobacteria, hydrogenases are suggested to drive the electron transport chain in the absence of exogenous electron acceptors (Berney and Cook, 2010), allowing continued production of ATP.

While menaquinones are the main lipoquinone for mycobacteria during aerobic growth, the biosynthesis of isoprenoid precursors is markedly downregulated during hypoxia, resulting in a depletion of menaquinones (Honaker et al., 2010; Matsoso et al., 2005). Under such hypoxic conditions, addition of the MK-9 analogue MK-4 (vitamin K2) or the saturated form (vitamin K1) is harmful and reduces the survival of *M. tuberculosis* (Honaker et al., 2010). A more recent study demonstrated that hypoxic conditions in a biofilm lead to the biosynthesis of polyketide quinones, which are alternative electron carriers that are produced by the type III polyketide synthases (Anand et al., 2015). These previous studies indicate that lipoquinone biosynthesis is a highly regulated process, controlled by sensing changing environmental factors. Our study showed that a partial depletion of MenG leads to the accumulation of DMK-9 without significant changes in the MK-9 pool. We speculate that this imbalance of DMK-9 and MK-9, induced by the MenG depletion, has a global impact on metabolic activity. While further studies are needed to understand the complex changes in menaquinone metabolism during MenG depletion, our current study highlights the spatial complexity of menaquinone biosynthesis, and the essential role of MenG, an IMD-associated protein, in maintaining the metabolic homeostasis and the active growth of *M. smegmatis*.

## Acknowledgements

This work was supported by grants from the Pittsfield Anti-Tuberculosis Association to YSM and the NIAID to DBM (AI 111224, AI 049313), and the University of Massachusetts Graduate School Dissertation Research Grant to JP. JP is a recipient of the Science Without Boarders Fellowship from CAPES-Brazil (0328-13-8). We thank Dr. Stephen Eyles (the Mass Spectrometry Center, the Institute of Applied Life Sciences, University of Massachusetts Amherst) for help with mass spectrometry.

